# Fine-tuning Nanog expression heterogeneity by altering MicroRNA regulation

**DOI:** 10.1101/2020.08.28.273177

**Authors:** Tagari Samanta, Sandip Kar

## Abstract

Nanog exhibits a robust heterogeneous expression pattern within a population of embryonic stem cells (ESCs) under varied culture conditions. However, an efficient way to fine-tune the Nanog expression heterogeneity remains elusive. Herein, by employing a stochastic modeling approach, we demonstrate that Nanog expression heterogeneity can be controlled by modulating the regulatory features of a Nanog transcript specific microRNA. We demonstrate how and why the extent of origin dependent fluctuations in Nanog expression level can be altered by varying either the binding efficiency of the microRNA-mRNA complex or the expression level of the respective microRNA. Moreover, our model makes experimentally feasible and insightful predictions to maneuver the Nanog expression heterogeneity explicitly to achieve cell-type specific differentiation of ESCs.

## Introduction

Nanog is an important transcription factor that regulates the expression level of several cell fate-determining genes to uphold the pluripotent state of embryonic stem cells (ESCs) [1,2]. In principle, by altering the expression level of Nanog, one can systematically maneuver the proliferation and differentiation propensities of ESCs under specific growth conditions [3–5]. However, in ESCs, Nanog gets expressed in a highly heterogeneous manner under various culture conditions [6]. It has been argued that this heterogeneity in Nanog expression stems from fluctuations originated due to intrinsic (due to gene expression variability or molecular noise) and extrinsic (cell-to-cell variability) noise sources present in and around the ESC culture medium [7–9]. Moreover, experiments suggest that the overall fluctuations at the Nanog transcriptional level, independent of culture conditions, get robustly partitioned into intrinsic and extrinsic noises by maintaining a ratio of 45:55 [10]. This indicates that to modify Nanog level in a specific way, in the first place, we need to have a thorough and quantitative understanding of how to fine-tune the existing heterogeneity in Nanog expression within an ESC culture condition. Unfortunately, our knowledge in this direction is still very limited. However, developing possible means to control the Nanog expression variability can potentially lead to novel stem cell-based therapies in the future.

Recently, by proposing a stochastic numerical framework, we demonstrated that under the different culture conditions (with only serum, or in presence of inhibitors such as PD0325901, CHIR99021 and the different stoichiometric mixture of these two inhibitors (2i condition)), the ESCs retain the robust Nanog expression heterogeneity by dynamically adjusting the events associated with Nanog transcription [11]. To be specific, we showed that in various culture conditions, the rates related to the transcription of Nanog gene and the corresponding translational rates are altered in such a manner that a strict ratio of intrinsic and extrinsic fluctuations is maintained. Moreover, our model predicted that by introducing an excess of CHIR99021 (a GSK3 inhibitor) in the culture medium, the noise partitioning ratio can be shifted from 45:55 value. However, having an excess of CHIR99021 inhibitor can be quite toxic to the cells. Thus, we need to figure out an alternative and biologically feasible route to control the existing heterogeneity in Nanog transcription. In this regard, utilizing microRNA (miR) mediated transcriptional regulation can be an effective option to adjust the Nanog expression heterogeneity.

MicroRNAs are short nucleotide non-coding RNAs (~ 22 nt) that post-transcriptionally regulate gene expression by destabilizing the target mRNAs and reduce the translation efficiency of the target mRNAs [12,13]. Thus, miRs seem to play an important role in many biological processes such as tumor suppression [14], animal development [15], programmed cell death [16,17], synaptic development [18,19] and hematopoietic cell fate decisions [20,21]. The functional impact of miR binding depends on the complementarity between miR and their mRNA targets. Recent studies have confirmed that by creating an imperfect base pairing between miR and the target mRNA, the translation repression can be impacted without affecting the target mRNA degradation [22,23]. Whereas, perfect or nearly perfect homology leads to faster mRNA degradation via endonucleolytic cleavage [24–26]. Intriguingly, miRs, in general, prefer to target genes with low expression, and eventually decrease the fluctuations observed for such genes. This advocates that varying miR binding efficiency with target mRNA, or by modifying the expression level of miR, one can influence the variability observed for the target gene expression. It has also been established in the literature that miRs can reduce the fluctuations at the protein level of the corresponding target gene in a context-dependent manner, and variability in their expression can further impart additional noise in the biological system [27].

Importantly, there exists a specific microRNA (miR-296), which specifically targets mRNA of Nanog [28], and regulates the transcription and translational efficiency of Nanog transcript significantly. How it affects the expression heterogeneity of Nanog, however, remains unexplored. In this article, we unfold the effect of miR-296 on Nanog expression heterogeneity by modifying our previously proposed model for the bi-allelic Nanog transcriptional reporter system. We systematically introduce the effect of miR-296 in Nanog transcriptional network and quantify the contributions of intrinsic and extrinsic fluctuations in Nanog expression using our previously proposed algorithm. Our study demonstrates how Nanog expression heterogeneity and specifically the ratio of intrinsic and extrinsic fluctuations can be influenced by simply modifying either the miR-296 and Nanog mRNA binding efficiency, or the expression level of miR-296 itself. Analysis of our stochastic simulations qualitatively explains the miR-296 mediated noisy regulation of Nanog expression and provides further insights about how to modify the pluripotent state of ESCs systematically.

## The Model

Over the last few decades, several network models with increasing complexities had been proposed [29–32] to decipher the underlying network dynamics that control the core transcriptional regulation in ESCs by explaining the state of the art experimental data. Even the heterogeneous Nanog gene expression and reprogramming efficiency had been analyzed with great details by considering the stochastic modeling setup [6]. This model captures the intrinsic variability of the corresponding network by implementing Gillespie’s stochastic simulation algorithm (SSA) [33], but cannot account for various sources of extrinsic fluctuations that turned out to be crucial to understand Nanog transcriptional heterogeneity [34]. However, recently, we proposed a simple core transcriptional regulatory network of Nanog that can efficiently measure the relative proportions of intrinsic and extrinsic fluctuation observed in Nanog transcript and protein dynamics by explaining a range of experimental observations [11]. The algorithm that we proposed based on SSA, not only allows us to quantify the heterogeneity in Nanog dynamics arising due to molecular fluctuations (intrinsic noise) of the species present in the respective network, but it further estimates the noise that originates due to fluctuations other than the molecular fluctuations (extrinsic noise). In this regard, our proposed method was unique and advantageous to quantify fluctuations that originate out of a bidirectional transcriptional reporter system (like in this case, MS2 and PP7 transcripts [10,35] or mRNAs of same Nanog protein [36]), where the transcription process remains under the control of the same promoter, while the corresponding gene gets simultaneously transcribed from two different transcripts (or mRNA’s) at the same time. Wherein, we were able to reconcile varied experimental findings [10,37–40] that deal with the amount of intrinsic and extrinsic fluctuations in Nanog regulation. Importantly, the same method can be applied for other genetic networks as well, so it is a general method.

In the current Nanog transcriptional network (**Fig. 1A** and **SFig. 1**), we made a considerable amount of modifications to our previously proposed Nanog transcriptional regulatory network [11] to study the effect of miR-296 over Nanog expression heterogeneity, both at the mRNA and protein level. The model proposed earlier modeled the bi-directional transcriptional activation of Nanog with simple mass-action kinetic equations, having the positive feedback regulation of Nanog imparted via activation of Oct4 and Sox2 proteins. The effect of the inhibitors to this transcriptional network is mostly modeled phenomenologically to reproduce the experimentally observed heterogeneity in Nanog transcriptional regulation. However, our previous model of core Nanog transcriptional regulation lacks various important interactions (such as Nanog autorepression [41], the repressive effect of Oct4/Sox2 at high Oct4 concentration [42] and related Nanog and Oct4/Sox2 autoregulation [4,32] that are known experimentally.

**Fig. 1.**
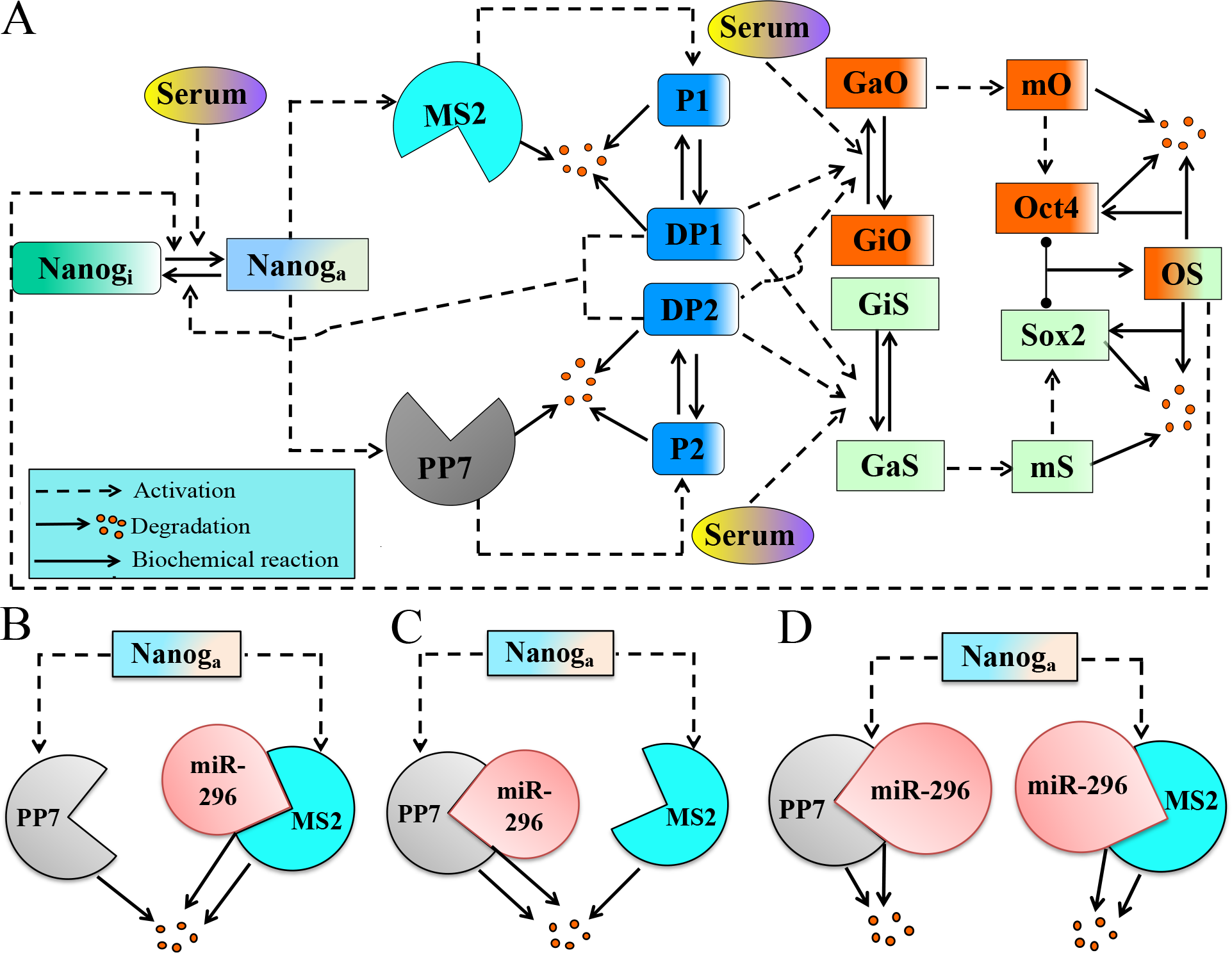
The proposed network of Nanog regulation under the influence of miR-296. **(A)** The detailed network of Nanog regulation that includes the two Nanog transcripts (MS2 and PP7), the Nanog protein (P1 and P2), the dimeric forms of Nanog protein (DP1 and DP2), and the crucial positive feedback interaction via Oct4 and Sox2 through the corresponding Oct4 and Sox4 heterodimer (OS). In our study, we consider that the miR-296 can influence the Nanog transcript in the following ways; **(B)** miR-296 specifically binds to the 3’UTR of MS2 transcript, **(C)** miR-296 specifically binds to the 3’UTR of PP7 transcript, and **(D)** miR-296 can bind with both the Nanog transcripts (MS2 and PP7) in a non-specific manner.

To begin with, we introduced those important additional interactions in our newly proposed network (**Fig. 1A** and **SFig. 1**) and modified our model equations (**Table-S1**) to include these effects accordingly. In this way, our Nanog transcriptional regulatory network now contains all the known regulatory nodes and is equipped with measuring the heterogeneity arising due to both intrinsic and extrinsic fluctuations efficiently for the corresponding network at the mRNA level by mimicking the smFISH kind of experiments [10] possible due to the presence of two Nanog transcripts (MS2 and PP7) in the network. However, with our previous model, computing the fluctuations at the protein level by following the same protocol [11] was not possible, as we assumed that both the MS2 and PP7 transcripts of *Nanog* genes are producing the same Nanog protein. In the current network (**Fig. 1A**), we proposed that both the transcripts can produce functionally similar Nanog proteins (*P*_1_ and *P*_2_) with different fluorescently tagged probes [43]. Thus, by counting the number of Nanog proteins generated from MS2 and PP7 transcripts via single-cell level experiments, one can measure the fluctuations at the protein level. Since Nanog protein forms dimer before acting as a transcription factor, we introduced the corresponding dimers (*DP*_1_ and *DP*_2_)) of *P*_1_ and *P*_2_ in the model [44]. We further assume that both *DP*_1_ and *DP*_2_ are inhibiting the transcription process of Nanog directly, and the Oct4/Sox2 complex (*OS*) can inhibit the process of transcription as the level of Oct4 and consequently the *OS* complex goes above a certain level (**Table-S1**).

With this improved basic structure of the Nanog transcriptional network, we explore the effect of miR-296 on Nanog expression heterogeneity systematically and sequentially by further extending the network (**Fig. 1B-D**). With knowledge of binding sites present in the 3’untranslated region (3’UTR) of Nanog transcripts under consideration, experimentally, it is possible to construct specific forms of miR-296 that will exclusively bind to either MS2 (**Fig. 1B**) or PP7 (**Fig. 1C**) Nanog transcript. These specific kinds of miR-296 will only regulate the number of transcripts and the corresponding translational efficiency of the respective transcript for which it is specifically made. We further consider that even other types of miR-296 can be produced experimentally, which can non-specifically bind with both the MS2 and PP7 Nanog transcripts (**Fig. 1D**), and can control the expression and translational efficiency of both the transcripts simultaneously. The Nanog regulatory dynamics get even more complicated under various culture conditions that contain conventional inhibitors like PD0325901 (inhibits ERK signaling)[45–48] and CHIR99021 (inhibits GSK-3 signaling) [49–51]. In our earlier work, we showed how these two inhibitors individually and collectively (2i condition) govern the Nanog transcriptional heterogeneity [11]. It turns out that these two inhibitors, in general, even regulate the miR processing significantly by influencing the two important miR processors DGCR8 and Drosha, respectively. Recent reports suggest that MEK/ERK can phosphorylate DGCR8, which increases its intracellular stability and hence induces a pro-growth miR profile [52,53]. On the other hand, GSK3 phosphorylates the Drosha and increases its nuclear localization [54–56]. Thus, phosphorylation of DGCR8 and Drosha gets repressed by PD0325901 (MEK inhibitor) and CHIR99021 (GSK3 inhibitor), which could result in the loss of miR expression. In our proposed model (**Fig. 1** and **SFig. 1**), we have included these effects of inhibitors in miR-296 expression phenomenologically, as very little is known about how exactly these inhibitors are functioning to suppress the expression level of miR-296 (**see supplementary file**).

Based on these facts, we have developed the deterministic ordinary differential equation model for the whole network proposed in **Fig. 1** for different miR-296 interactions (**Fig. 1B-D**), and it is provided in **Table-S1**. We have a detailed account of each mathematical terms included in the model in the supplementary section that follows **Table-S1**. The chemical reactions (**Table-S2**) involved in the network are simulated by following the stochastic simulation algorithm to quantify both the intrinsic and extrinsic fluctuations. The abbreviation of the respective variables used while constructing the ordinary differential model, and the parameter values are described in **Table-S3** and **Table-S4**, respectively. In **Table-S4**, we have mentioned, which parameters are known from experiments or are earlier used in the literature. We have further indicated the other parameters that are chosen to reconcile the experimental observations related to Nanog heterogeneity. The related model ODE file and the stochastic code are freely available in the Github repository (https://github.com/tagarisamanta/Nanog_microRNA.git).

## Method

We have constructed the deterministic part of the model by using mostly mass-action kinetic terms and added the effect of the inhibitors [45, 47–53] later by introducing phenomenological kinetic terms [11]. This allowed us to reproduce the average number of mRNA and protein expression levels, as observed in the single-cell experiments [10] (**See supplementary section-1, for details**). Furthermore, we employed our recently proposed stochastic simulation algorithm (**SFig. 2**) that we developed based on Gillespie’s stochastic simulation algorithm (SSA)[33]. This newly devised stochastic simulation method allows us to quantify the intrinsic and extrinsic fluctuation adequately for our proposed Nanog transcriptional network (**Fig. 1**). The overall stochastic algorithm is described in detail in **SFig. 2,** and more details about the method can be obtained from our previous paper [11]. Our stochastic simulation method accounts for both the intrinsic noise (arises mainly due to gene expression variability, commonly termed as molecular fluctuation) and the extrinsic fluctuation (sources of extrinsic variability can be of different origin, for example; (i) cell division process [34], (ii) the changes in the transcriptional efficiency during different phases of the cell cycle [57] (iii) the varied initial condition of different species related to the transcriptional network [58,59], etc..) in a much efficient manner by taking care of the various additional noise sources other than the molecular fluctuation, which is inherent to any biological network. For further details about the stochastic simulation method, we refer to the method section of our previous paper [11] and discussion made therein (and **SFig.2**), where a detailed account for the model implementation for the stochastic simulation algorithm had been laid out explicitly, which we followed in this manuscript as well. In each case, we performed our stochastic simulation for about 2000 cells (in the context of simulation, each ES cell is a unique random no seed used for that stochastic run) to report our results.

## Results and discussion

### Altering binding efficiency of transcript specific miR-296 disparately fine-tunes Nanog expression variability

We initiated our work by hypothesizing that the miR-296 is specifically designed to bind with a specific Nanog transcript i.e., either the MS2 or PP7 probe containing Nanog transcripts. In recent literature [27], there are examples of constructing such transcript specific microRNA binding situation by genetically modifying the 3’UTR region of transcript itself, such that it contains one or two specific microRNA binding site(s). Analyzing this scenario seems quite relevant, as the Nanog transcripts with MS2 and PP7 probes do have different reported half-lives. Since the stability of a transcript critically determines the level of intrinsic fluctuations, we speculated that a transcript specific miR-296 **(Fig. 1B)** might affect the robustness inherent to the Nanog transcriptional heterogeneity. Indeed, we made pretty interesting observations. In our simulations, we started with a specific miR-296-MS2-transcript interaction (*k*_mir1_) rate (where a ~45:55 ratio of intrinsic:extrinsic is maintained) and systematically increase the same, while measuring the different kinds of fluctuations as a function of total mRNA number. Initially, we detect a steady decrease and increase of intrinsic (**Fig. 2A, left panel**) and extrinsic (**Fig. 2A, middle panel**) fluctuations, respectively, as a function of *k*_mir1_. Fluctuations at the Nanog protein level (**SFig. 3**) follow a similar pattern, which is following the recent experimental observation made by Schmiedel et al. [27] in a similar context. A further rise in *k*_mir1_, however, leads to an increase in intrinsic fluctuation, and the same trend gets reflected in the total noise variability (**Fig. 2A, right panel**). Consequently, the ratio of intrinsic and extrinsic fluctuations deviates significantly from 45:55 value (**Fig. 2B** and **SFig. 4**) due to the higher contribution of extrinsic variability induced by altering the miR-296-MS2-transcript interaction under varied specific total Nanog mRNA levels. Interestingly, numerically introducing a PP7 transcript specific miR-296 in the network (**Fig. 1C**), systematically increases both the intrinsic (**Fig. 2C, left panel**) and extrinsic (**Fig. 2A, middle panel**) fluctuations as a function of *k*_mir2_, where the total noise (**Fig. 2A, right panel**) variation pattern is completely dictated by the intrinsic fluctuation pattern. **Fig. 2D** suggests that the ratio of intrinsic and extrinsic noises can be shifted from 45:55 value by modifying the miR-296-PP7-transcript interaction, however, in this case, intrinsic fluctuation impacts the ratio to a greater extent.

**Fig. 2.**
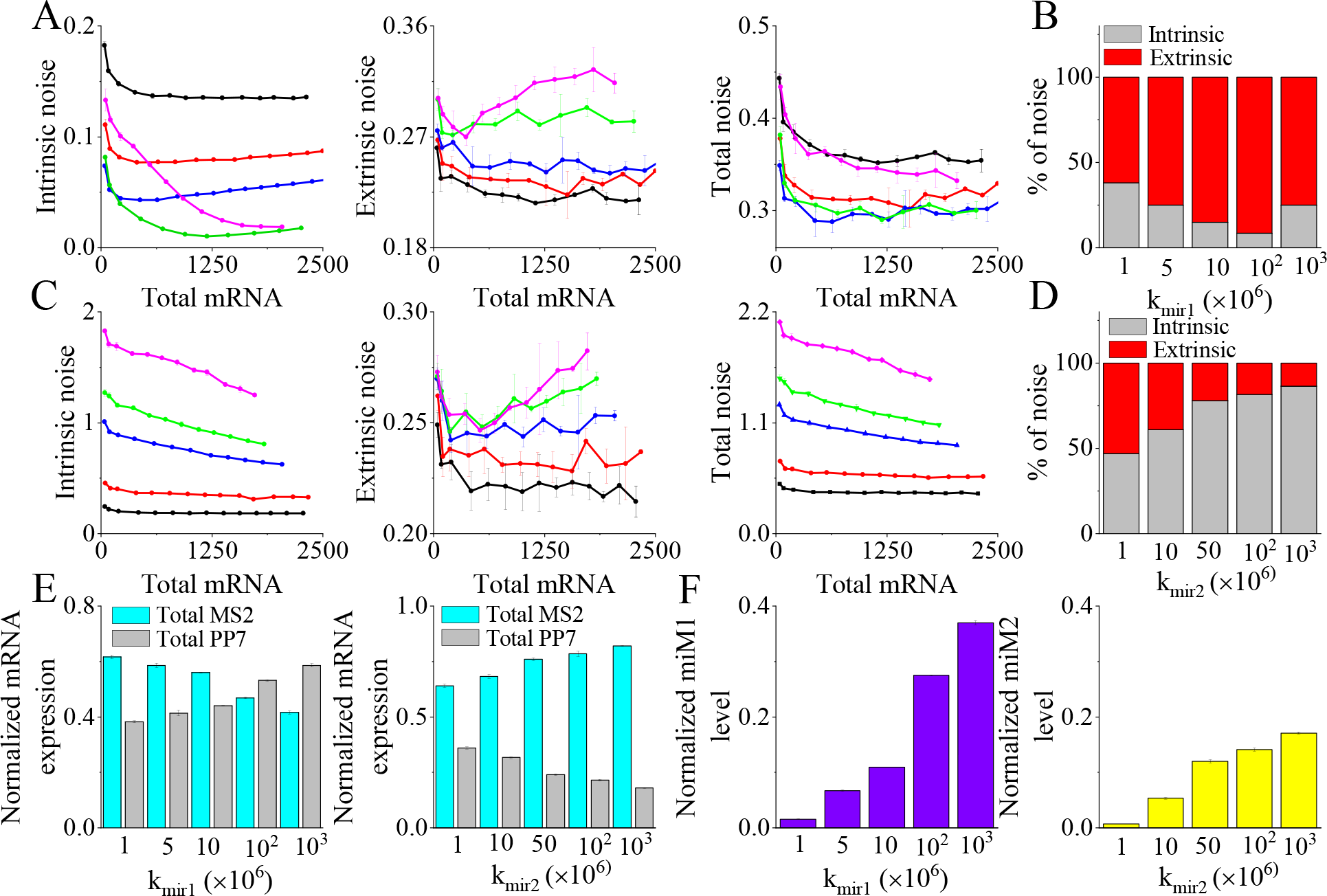
Altering binding efficiency of transcript specific miR-296 disparately fine-tunes Nanog expression variability. **(A)** Various fluctuations (intrinsic (left panel), extrinsic (middle panel), and total noise (right panel)) in Nanog mRNA are quantified in presence of MS2 transcript specific miR-296 for different miR-296-MS2-transcript binding efficiencies (k_mir1_ = 10^−6^ (Black-WT), 5×10^−6^ (Red), 10^−5^ (Blue), 10^−4^ (Green) and 10^−3^ (Pink), all in molecule^−1^min^−1^) as a function of total mRNA number. **(B)** The percent contribution of intrinsic and extrinsic noise is plotted for different miR296-MS2-transcript binding rates. **(C)** Various fluctuations (intrinsic (left panel), extrinsic (middle panel), and total noise (right panel)) in Nanog mRNA are quantified in presence of PP7 transcript specific miR-296 for different miR-296-PP7-transcript binding efficiencies (k_mir2_ = 10^−6^ (Black-WT), 10^−5^ (Red), 5×10^−5^ (Blue), 10^−4^ (Green) and 10^−3^ (Pink), all in molecule^−1^min^−1^) as a function of total mRNA number. **(D)** The percent contribution of intrinsic and extrinsic noise is plotted for different miR296-PP7-transcript binding rates. **(E)** Normalized mRNA expression (free + complexed) of MS2 and PP7 transcripts in presence of various MS2 specific (left panel) and PP7 specific (right panel) miR-296-Nanog-transcript binding rates. **(F)** The normalized (concerning the corresponding total mRNA expression) count of miR-296-MS2-transcript (left panel) and miR-296-PP7-transcript (right panel) complexes to the total Nanog mRNA expression is plotted under the different extent of respective miR-296-Nanog-transcript interaction.

To interpret these interesting observations (**Fig. 2A-D**), in **Fig. 2E** and **Fig. 2F**, we compared the respective measures that allowed us to compare the intrinsic and extrinsic fluctuations under different conditions. Experimentally, it has been demonstrated that the Nanog MS2 transcript has higher stability than the corresponding PP7 transcript, and under wild-type (WT) situation (**Table-S4**), the number of MS2 transcript is more than PP7 transcript. Thus, introducing an MS2 transcript specific miR-296 (keeping all other interactions the same) will eventually reduce (**Fig. 2E, left panel**) the difference between the expression levels of MS2 and PP7 transcripts, as the miR-296-MS2-transcript interaction is increased. This results in the initial sharp decrease of intrinsic fluctuation under these conditions. However, a further rise in the miR-296-MS2-transcript interaction leads to an increase in the intrinsic fluctuation as the difference between the expression levels of MS2 and PP7 again goes up (**Fig. 2E, left panel**). Conversely, a PP7 transcript specific miR-296 monotonically increases the difference of the expression levels of MS2 and PP7 transcripts (**Fig. 2E, right panel**), and promotes the extent of intrinsic fluctuations as a function of miR-296-PP7-transcript interaction.

**Fig. 2E** elucidates the trends in the intrinsic fluctuation under both the scenarios, but the extrinsic variability and the corresponding ratios of intrinsic and extrinsic fluctuations can not be solely explained by this analysis. The extent of extrinsic variations as a function mRNA-miR interaction can be quantified by measuring the level of the mRNA-miR complex under respective conditions. In **Fig. 2F**, it is evident that the normalized level of miR-296-MS2-transcript complex (miM1) grows relatively sharply (**Fig. 2F, left panel**), as the miR-296-MS2-transcript interaction is amplified in the numerical thought experiment. This eventually tilts the ratio of intrinsic and extrinsic fluctuations by rising the influence of extrinsic fluctuation (**Fig. 2B**). However, the relative rise in the normalized level of miR-296-PP7-transcript complex (miM2) is less steep (**Fig. 2F, right panel**) than the previous case, leading to a sluggish upsurge of the extrinsic noise. Thus, a combination of a substantial increase in the intrinsic noise supplemented with a slow increase in extrinsic noise alters the ratio of intrinsic and extrinsic fluctuations more in favor of intrinsic fluctuations.

### Nonspecific miR-296 binding makes Nanog expression heterogeneity more dependent on extrinsic fluctuations

In the previous section, we have shown that introducing transcript specific miR-296, one can systematically modify the Nanog fluctuation both at the mRNA and protein level. Here, we further explore the effect of nonspecific miR-296 regulation on Nanog expression heterogeneity. It is important to mention that creating a nonspecific miR-296 binding scenario would be relatively easier experimentally [27], as identical modifications to the 3’UTR region of both the transcripts will produce such nonspecific binding automatically. We assumed that the miR-296 will bind with both the MS2 and PP7 Nanog transcripts with equal efficiency. First, we have performed our stochastic simulation by considering that the degradation rates of the Nanog transcripts from the respective miR-296-mRNA complexes are ~2.5 times that of the normal degradation rates of the individual Nanog transcripts (**Table-S4**). Our model simulations reveal that under this condition, the intrinsic fluctuations remain almost unaltered as a function of miR-296-mRNA binding efficiencies (*k*_mir1_ and *k*_mir2_), and decreases monotonically with the increased expression level of total mRNA under any specific value of miR-296-mRNA binding rate (**Fig. 3A, left panel**). This observation suggests that the relative counts of the MS2 and PP7 transcripts should remain almost constant as the miR-296-mRNA binding rates are varied, and **Fig. 3B** displays the same.

**Fig. 3.**
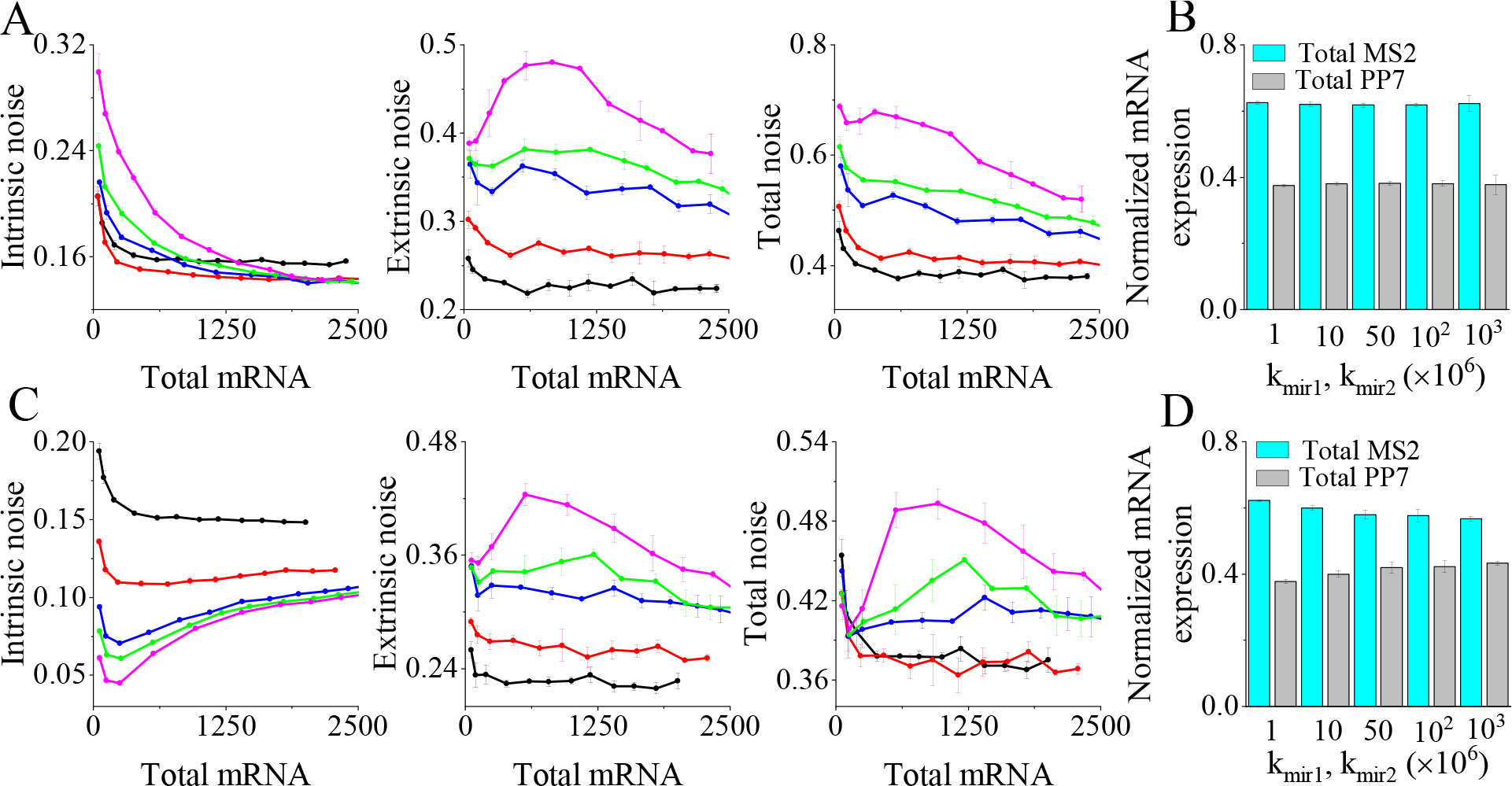
Understanding Nanog transcriptional heterogeneity in the presence of non-specific miR-296 binding with both the Nanog transcripts. **(A)** Various fluctuations (intrinsic (left panel), extrinsic (middle panel), and total noise (right panel)) in Nanog mRNA are quantified in presence of non-specific miR-296 binding with both MS2 and PP7 transcripts for different miR-296-Nanog-transcript binding efficiencies (k_mir1_ and k_mir2_ = 10^−6^ (Black-WT), 10^−5^ (Red), 5×10^−5^ (Blue), 10^−4^ (Green) and 10^−3^ (Pink), all in molecule^−1^min^−1^) as a function of total mRNA number. (here, the same ratio of degradation rates of mRNA from free and complexed forms (i.e., k_dmim1_: k_dmim2_ ≈ k_dm1_: k_dm2_) are considered). **(B)** The normalized expression levels of total MS2 and PP7 transcripts are plotted for different miR-296-Nanog-transcript binding rates considering the data used in **Fig. 3A**. The normalization is done concerning the corresponding total mRNA level. **(C)** Various fluctuations (intrinsic (left panel), extrinsic (middle panel), and total noise (right panel)) in Nanog mRNA are quantified in presence of non-specific miR-296 binding with both MS2 and PP7 transcripts for different miR-296-Nanog-transcript binding efficiencies (k_mir1_ and k_mir2_ = 10^−6^ (Black-WT), 10^−5^ (Red), 5×10^−5^ (Blue), 10^−4^ (Green) and 10^−3^ (Pink), all in molecule^−1^min^−1^) as a function of total mRNA number. In this case, we assume that k_dmim1_> k_dmim2_. **(D)** The normalized expression levels of total MS2 and PP7 transcripts are plotted for different miR-296-Nanog-transcript binding rates considering the data used in **Fig. 3C**. The normalization is done with respect to the corresponding total mRNA level.

On the contrary, the extent of extrinsic fluctuation rises quite steeply as miR-296-mRNA binding rates are increased (**Fig. 3A, middle panel**), and the total noise (**Fig. 3A, right panel**) in the Nanog transcription varies similarly. While the trend in extrinsic noise variation can be accounted for by a sharp rise in the relative level of total miR-296-mRNA complex (**SFig. 5A**), the greater dependence on extrinsic noise again predicts a noteworthy deviation (**SFig. 5B**) from the 45:55 (intrinsic: extrinsic) ratio normally associated with Nanog transcriptional regulation.

The insensitiveness of intrinsic fluctuation as a function of miR-296-mRNA binding rate might be due to our initial assumption, where we tacitly assume that the ratio of the degradation rates of the Nanog transcripts from the respective miR-296-mRNA complexes is same as that of the ratio of the individual normal degradation rates of the respective transcripts. This hypothesis ensures that the number of MS2 and PP7 transcript varies correspondingly, while the miR-296-mRNA binding rates are altered. Here, we kept the degradation rates of MS2 transcript (both from miR-296-mRNA complex and the individual transcript) the same as taken for **Fig. 3A**, but assumed that the PP7 degradation rate from respective miR-296-mRNA complex is marginally higher than its normal degradation rate. Our stochastic simulations under these parametric conditions depict that the intrinsic noise will decrease (**Fig. 3C, left panel**) as a function of miR-296-mRNA binding rate. This variation in intrinsic noise can again be explained qualitatively by measuring the total number of MS2 and PP7 transcripts (**Fig. 3D**) under such conditions. In this instance too, the overall noise (**Fig. 3C, right panel**) continues to be more dependent on the extrinsic variability (**Fig. 3C, middle panel**) with increasing complex formation rate between miR-296 and the Nanog transcripts. We further made two very interesting observations at higher values of miR-296-mRNA binding rate. The intrinsic fluctuation showed an increasing trend, while the total noise as well as the extrinsic fluctuation passed through a maximum with an increase in the total mRNA number at a higher miR-296-mRNA binding rate. We will discuss these observations in detail in the next section.

### The model predicts unusual trends in Nanog expression heterogeneity under miR-296 overexpression conditions

Till now, we have mostly concentrated on understanding the effect of binding efficiency of miR-296 and Nanog transcripts on Nanog heterogeneity. Nonetheless, experimentally, producing these diverse kinds of miR-296 binding scenarios with varying complementarity is a challenging task. However, under certain miR-mRNA binding rates, over-expressing miR-296 within an embryonic cell seems to be a relatively easier experiment to perform. Keeping this fact in mind, we performed stochastic simulations in different miR-296 over-expression conditions at relatively low and high miR-296-mRNA binding rates. Here, the degradation rates of the respective transcripts are taken as it was in the case of **Fig. 3C-D**. Our simulations show that at lower binding efficiency, intrinsic fluctuation reduces (**Fig. 4A, left panel**), as the expression level of miR-296 is increased gradually. Under the same situation, extrinsic noise remains mostly unaffected (**Fig. 4a, middle panel**), and total noise variation (**Fig. 4A, left panel**) follows the pattern of extrinsic noise variability. **SFig. 6** qualitatively rationalizes the abovementioned observations. A steady drop in the difference between the total average number of MS2 and PP7 transcripts (**SFig. 6A**) corroborates with the monotonic decrease in intrinsic fluctuation, while insignificant variation in the number of miR-mRNA complex (**SFig. 6B**) under different expression level of miR-296 justifies the extrinsic fluctuation pattern.

**Fig. 4.**
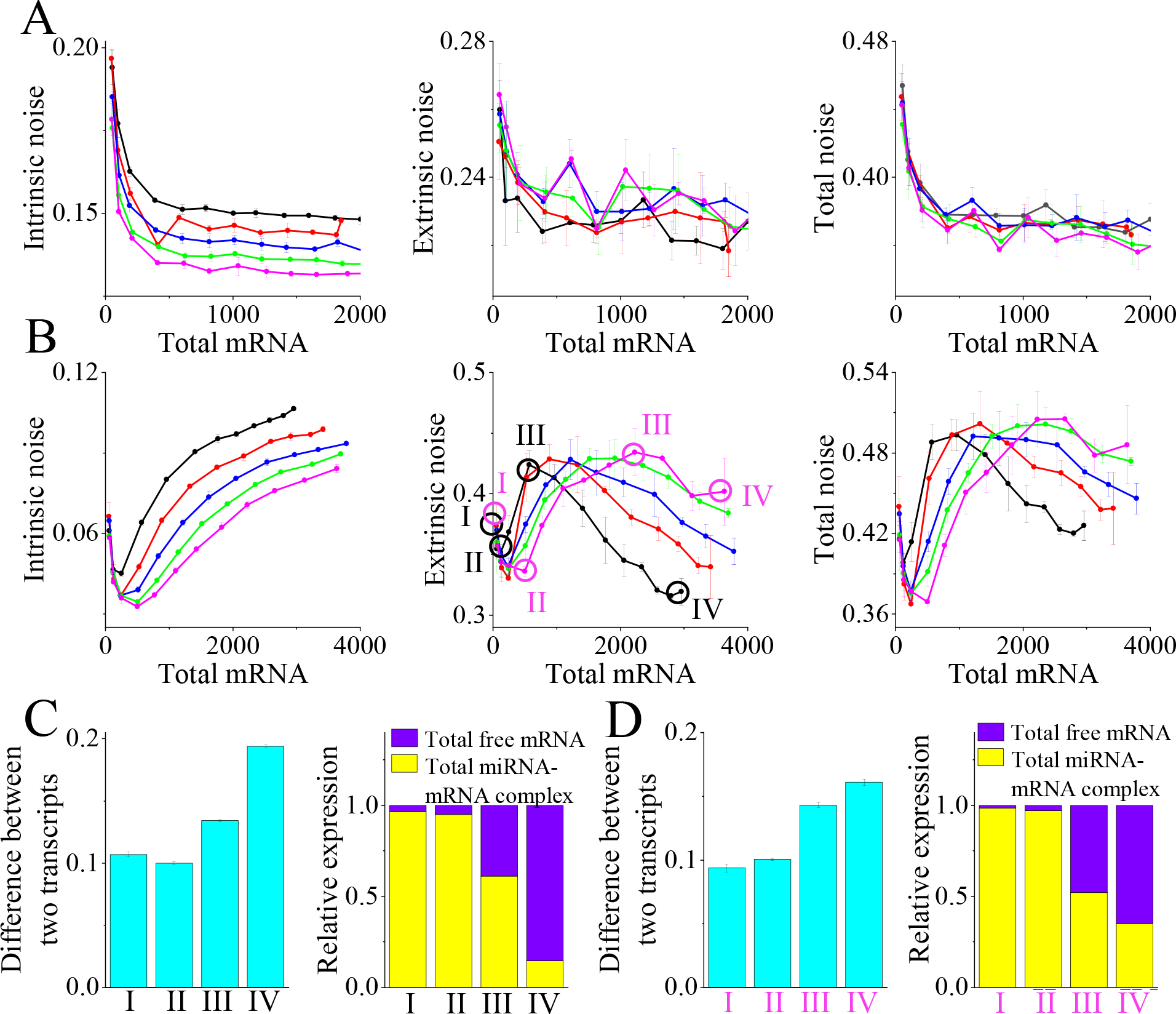
Deciphering the effect of miR-296 over-expression on Nanog expression heterogeneity. Quantifying the fluctuations (intrinsic (left panel), extrinsic (middle panel), and total noise (right panel)) under different overexpression levels (b_mir_ = 1 (black), 1.5 (red), 2 (blue), 2.5 (green) and 3 (pink) molecule min^−1^) of miR-296 for **(A)** k_mir1_, k_mir2_ = 10^−6^ molecule^−1^min^−1^ (WT situation), and **(B)** highest k_mir1_, k_mir2_=10^−3^ molecule^−1^min^−1^ values. The normalized difference between the total MS2 and PP7 transcripts (left panels) and the relative expression of the respective total free transcript to miR-296-Nanog transcripts (right panels) for various level of total Nanog mRNA are plotted for four different total mRNA expression with; **(C)** highest miRNA binding efficiency (k_mir1_, k_mir2_ =10^−3^ molecule^−1^min^−1^) and lower miRNA basal synthesis rate (b_mir_=1 molecule min^−1^), and **(D)** highest miRNA binding efficiency (k_mir1_=10^−3^ molecule^−1^min^−1^) and highest basal synthesis rate of miRNA (b_mir_=3 molecule min^−1^). (In both **(C)** and **(D)**, I, II, III, and IV denote the respective total expression levels of Nanog mRNA and the corresponding total miR-296-Nanog-transcript numbers).

However, at higher miR-296-mRNA binding efficiency, our stochastic simulations uncover a completely counterintuitive fluctuation pattern under miR-296 over-expression condition. The intrinsic fluctuation passes through a minimum (**Fig. 4B, left panel**) with an increasing level of total Nanog transcript under any specific miR-296 expression level. Although the extrinsic noise shows a similar initial dip, eventually it goes through a maximum (**Fig. 4B, middle panel**) under the same condition. Total noise variation (**Fig. 4B, left panel**) again shadows mostly extrinsic noise variability. Intriguingly, the relative difference in the total transcript numbers of the two individual Nanog transcripts both at lower (**Fig. 4C, left panel**) as well higher (**Fig. 4D, left panel**) expression level of miR-296 for four different total mRNA levels qualitatively explains why the intrinsic fluctuation goes through a minimum. It is evident from these plots that the relative difference in the transcript numbers too passes almost through a minimum under similar circumstances. However, rationalizing the variation in extrinsic fluctuation qualitatively seems to be a challenging task. To understand the extrinsic variability (**Fig. 4B, middle panel**), we tried to quantify the relative amount of the total miR-296-mRNA complex at various corresponding total mRNA levels for different expression levels of miR-296.

It turns out that when the total transcript level of Nanog is much lower than the total miR-296 number, almost all the Nanog transcripts remain in the form of miR-296-mRNA complex (**Fig. 4C-D, right panels,** first two bars), and the relative abundance of miR-296-mRNA complexes does not correlate with the extent of extrinsic fluctuation. Under such conditions, variation in intrinsic fluctuation mostly dictates the total noise (**Fig. 4B, left panel**). However, as the total mRNA number becomes higher relative to the miR-296 level, the number variation in miR-296-mRNA starts correlating with the extrinsic variability (**Fig. 4C-D, right panels,** last two bars) and explains the nature of extrinsic fluctuations as well. Interestingly, the maximum of extrinsic noise appears under any specific expression level of miR-296, when the relative amount of miR-296-mRNA complex represents about half (**Fig. 4C-D, right panels,** third bar) of the total Nanog transcript level. At a much higher level of total mRNA number, the number of miR-296-mRNA complex reaches a saturation level, and extrinsic fluctuation gradually decreases with an increasing number of total mRNA (**Fig. 4C-D, right panels,** fourth bar). Thus, our qualitative analysis provides simple measures to quantify and analyze the complex fluctuation patterns that can be realized in Nanog expression heterogeneity.

### Inhibitory culture conditions maintain robust Nanog expression heterogeneity by downregulating miR-296 expression

At this juncture, we further investigated how various culture conditions containing either only PD0325901 (MEK inhibitor) or a mixture (2i condition) of PD0325901 and CHIR99021 (GSK3 inhibitor) inhibitors maintain the robust Nanog expression heterogeneity in presence of miR-296 regulation. In literature, it is known that these inhibitors down-regulate the synthesis of miR-296 [53], and we have phenomenologically incorporated such kind of inhibition in our model (**Table-S1**). **Fig. 5A** illustrates how exactly the total miR-296 number gets down-regulated in the presence of inhibitory conditions in comparison to the WT case (**Table-S4**). We implemented stochastic simulations under these inhibitory conditions by maintaining a moderate miR-296 expression level, and keeping the degradation rates of Nanog transcript’s, as it was in the case of **Fig. 3C**. First, we performed our stochastic simulation for a moderate binding efficiency between miR-296 and the Nanog transcripts (WT like) and found that the extrinsic fluctuation falls off steadily as a function of total mRNA number both under only PD0325901 (**Fig. 5B, left panel**) and 2i inhibitory conditions (**Fig. 5B, right panel**).

**Fig. 5.**
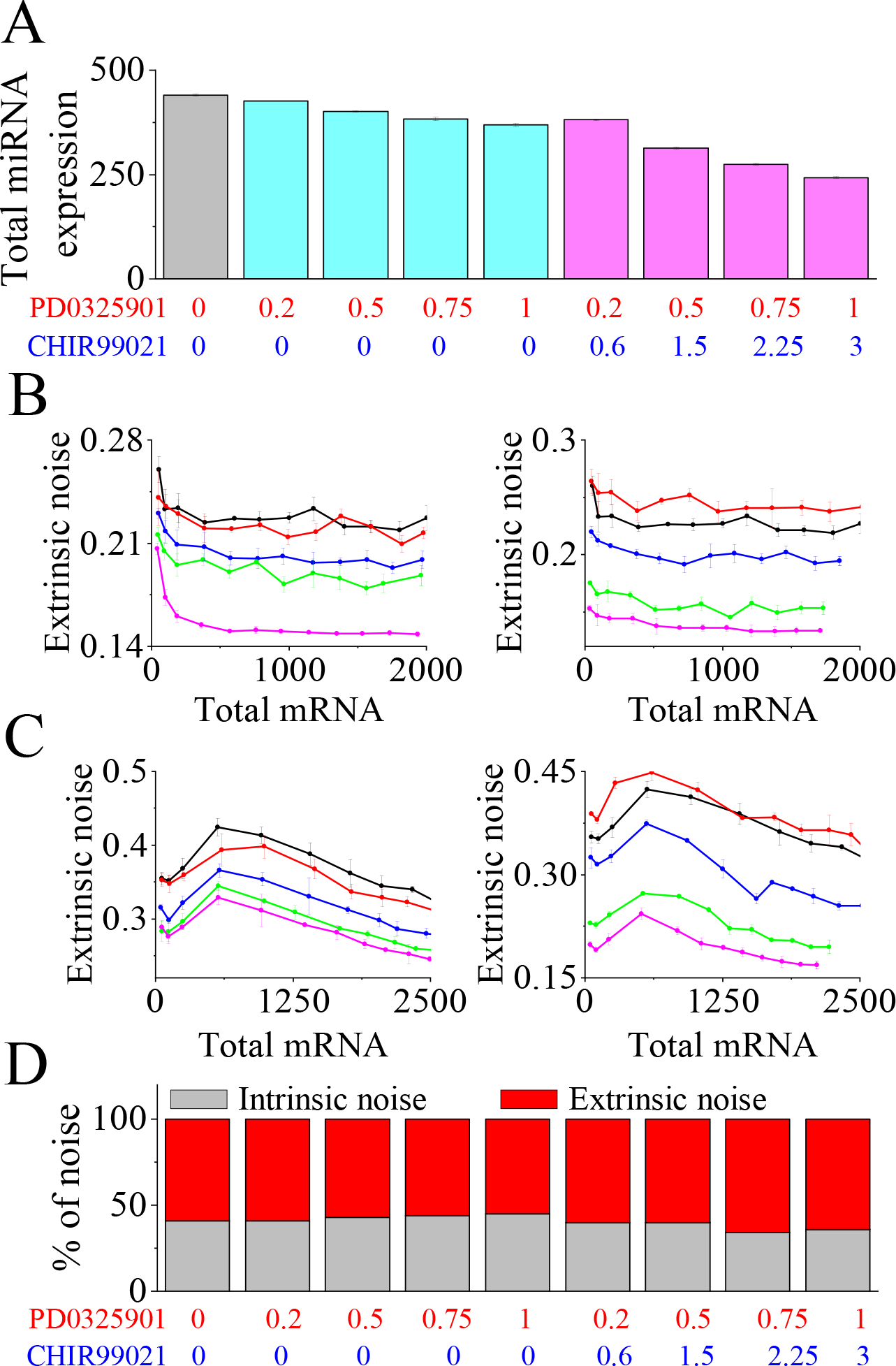
Effect of miR-296 on Nanog expression heterogeneity under different inhibitory culture conditions. (A) The variation of total miR-296 expression is shown under different inhibitory culture conditions. Extrinsic noise variation under different concentrations of only PD0325901 inhibitor (left panel) (I_1_= 0(black), 0.2(red), 0.5(blue), 0.75(green) and 1(pink)) and 2i conditions (right panel) (I_1_:I_2_= 0:0(black), 0.2:0.6(red), 0.5:1.5(blue), 0.75:2.25(green) and 1:3(pink)) with; **(B)** k_mir1_, k_mir2_ = 10^−6^ molecule^−1^min^−1^ and b_mir_ = 1 molecule min^−1^ (WT condition), and (C) k_mir1_, k_mir2_ = 10^−3^ molecule^−1^min^−1^ and b_mir_ = 1 molecule min^−1^. **(D)** The relative percentage of intrinsic and extrinsic fluctuations is plotted under different culture conditions for WT condition.

The same trends in extrinsic fluctuation (**Fig. 5C**) are found even at a higher binding rate of miR-296 and Nanog mRNAs. The decrease in extrinsic noise is mainly caused by the reduction in the miR-296 level under inhibitory conditions. However, the ratio of intrinsic:extrinsic fluctuations in Nanog transcription is strictly maintained around 45:55 when the miR-296-mRNA binding rate is kept at a moderate level (**Fig. 5D**). However, with increasing binding efficiency of miR-296 and Nanog transcripts, the same ratio differs significantly (**SFig. 7**) from conventionally observed 45:55 value. Thus, our model predicts diverse alternative paths to fine-tune Nanog expression heterogeneity in ESCs by simply altering miR-296 regulation.

### Sensitivity analysis of model parameters reveals ways to fine-tune the Nanog heterogeneity

Can we predict further avenues to alter the Nanog heterogeneity plausibly? To answer this important question, we have performed a systematic sensitivity analysis of our model parameters by taking the intrinsic, extrinsic and the total fluctuations observed at the Nanog mRNA level under normal WT type conditions as the sensitivity criteria. Sensitivity analysis (under low total transcript level) predicts (**Fig. 6A**) that intrinsic fluctuation at the Nanog transcript level critically depends on the respective degradation rates (*k*_dm1_ and *k*_dm2_) and the transcriptional frequencies (*k*_m1_ and *k*_m2_) of both of the Nanog transcripts (MS2 and PP7). However, the extrinsic variability at the Nanog mRNA level mostly remains sensitive to the parameters (*k*_ag_ and *k*_ag1_) governing the feedback of Oct4/Sox2 (OS) complex to the Nanog transcription (**Fig. 6B**). Interestingly, the sensitivity analysis taking the overall noise as the sensitivity criteria (**Fig. 6C**) demonstrates that the parameters, which are governing the intrinsic variability for the Nanog transcription, are even dictating the overall Nanog heterogeneity at the Nanog mRNA level. We have performed the sensitivity analysis with a high total transcript number of Nanog as well, and we found a similar trend of parameter sensitivity (**SFig. 8A**) for the model parameters.

**Fig. 6.**
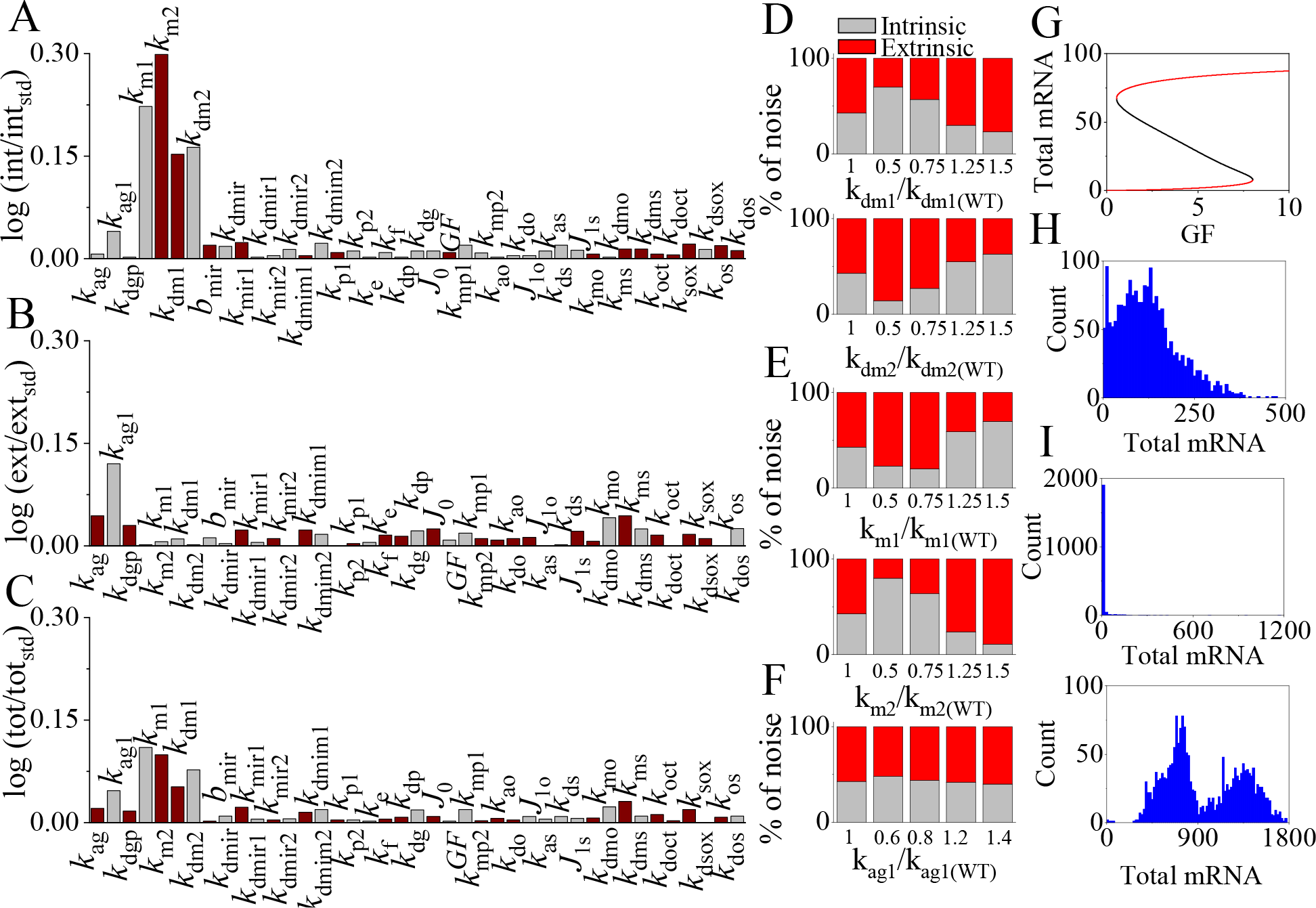
Sensitivity analysis of the model parameters predicts possibilities to fin-tune the Nanog heterogeneity in a systematic manner. Sensitivity analysis of the parameters involved in the model under normal growth condition by taking **(A)** intrinsic, **(B)** extrinsic, and **(C)** total noise as the sensitivity criteria. Kinetic parameters are increased individually (about 20% of the values provided in **Table-S4**) keeping all other parameters constant, and the relative increase or decrease in related noise criteria are denoted by grey and brown bars, respectively. Deviation from intrinsic:extrinsic noise ratio of 45:55 under WT condition by varying the: **(D)** degradation rates of MS2 (*k*_dm1_, **upper panel**) and PP7 (*k*_dm2_, **lower panel**) transcripts, **(E)** transcription rates of MS2 (*k*_m1_, **upper panel**) and PP7 (*k*_m2_, **lower panel**) transcripts, and **(F)** feedback related phenomenological parameter *k*_ag1_. **(G)** Bifurcation analysis of our proposed network shows bi-stability in Nanog dynamics. Parameters are as given in Table-S4, only *k*_ag_ is increased by 1.84 times, and subsequently to keep the GF regime the same, *J*_0_, *J*_10_ and *J*_1S_ are decreased by 30 times. **(H)** Stochastic simulation at GF=1 demonstrates a bi-modal Nanog distribution with both Nanog low and Nanog high cells. **(I) ES** Cells with low and high Nanog initial conditions are given 2i conditions, and low Nanog cells remained in the low Nanog state (**upper panel**), but cells with high Nanog maintained the high Nanog state (**lower panel**).

Experimentally, it is quite feasible to vary the degradation rates of the Nanog transcripts by genetically modifying them either individually or both at once [10]. Thus, at first, we altered the degradation rate of the Nanog transcript individually in our model, and our stochastic simulation revealed that making the MS2 transcript more stable (i.e., decreasing *k*_dm1_ concerning the WT case given in **Table-S4**) will increase the amount of intrinsic fluctuation (**Fig. 6D**, **upper panel**) over the extrinsic variation seen at the Nanog mRNA level. However, increasing *k*_dm1_ eventually raises the level of extrinsic variability substantially, and the overall Nanog heterogeneity depends overwhelmingly on the extrinsic fluctuation (**Fig. 6D**, **upper panel**). On the contrary, decreasing or increasing the degradation rate (*k*_dm2_) of the PP7 transcript of Nanog just gives a reverse effect (**Fig. 6D**, **lower panel**). Interestingly, changing both the degradation rates by maintaining the ratio of them intact (as it was in case of WT (**Table-S4**)), preserves the ratio of intrinsic:extrinsic with the normally reported ratio of 45:55 (**SFig. 8B**).

Altering transcription rates (*k*_m1_ and *k*_m2_) seem to be a much more challenging task experimentally. Nonetheless, we performed stochastic simulations by varying these two parameters individually and found that both *k*_m1_ (**Fig. 6E**, **upper panel**) and *k*_m2_ (**Fig. 6E**, **lower panel**) affect the Nanog heterogeneity quite appreciably, if decreased or increased concerning the WT values as given in **Table-S4**. It is also important to note that the phenomenological parameter *k*_ag1_ remains as one of the crucial regulators of the overall noisy regulation of Nanog transcription, as it shows up to be one of the parameters that can alter both intrinsic and extrinsic variability (**Fig. 6A-C**) to a noticeable extent. However, stochastic simulations by increasing or decreasing *k*_ag1_ individually (**Fig. 6F**) did not notably affect the Nanog heterogeneity. This is due to the fact that an increase or decrease in *k*_ag1_ affects both intrinsic and extrinsic variability in a similar manner (**Fig. 6F**), leading to the maintenance of the ratio of intrinsic:extrinsic intact in the range of 45:55.

However, the parameters *k*_ag1_ and *k*_ag_ are found to be the two most influential parameters that regulate the extrinsic fluctuation related to Nanog transcription (**Fig. 6B**). Importantly, the parameter *k*_ag_ determines the extent of both the positive and negative feedback regulation provided by the Oct4/Sox2 (OS) complex to the Nanog transcription. Wherein, in our model *k*_ag1_ sets the threshold for the concentration of OS, beyond which OS will give negative feedback to the Nanog transcription instead of positively regulating the same. To this end, with our current model, we performed a bifurcation analysis and found that by just increasing the *k*_ag_ parameter (i.e., by increasing the feedback regulation by OS), one can realize the bi-stable dynamics of Nanog (**Fig. 6G**), where a low Nanog level designates the differentiated state and high Nanog state stands for the pluripotent state of ES cells [60]. Stochastic simulation performed at GF=1 (**Fig. 6H**) demonstrates that Nanog mRNA distribution for a population of cells exhibits a bimodality, which is in agreement with the bi-stable dynamics of Nanog. We further challenged our model under this condition with the recent experimental findings by Hastreiter et al. [60], where they observed that in presence of 2i condition, Nanog maintains a high expression in ESCs, whereas, in Nanog low expressing cells, Nanog expression did not upregulate in presence of 2i condition. We implemented stochastic simulations by implementing 2i conditions (at GF=1) for Nanog low and Nanog high cells from the simulation performed only under GF=1 condition. Our stochastic simulation corroborates with the experimental findings and displays that Nanog low cells indeed maintains a Nanog low state (**Fig. 6I, upper panel**), whereas Nanog high cells uphold a Nanog high state (**Fig. 6I, lower panel**) even under 2i condition. Herein, we made a quick sensitivity analysis (**SFig. 9**) concerning the 2i condition related phenomenological parameters and found that the simulation results remained quite robust even if we changed those parameters to a reasonable extent. Thus, our model not only validates interesting experimental observations but reliably predicts varied ways to maneuver Nanog heterogeneity at the population level.

## Conclusion

Stem cell-based therapies certainly have great potential to deliver potent treatment strategies for various diseases in the future. Unfortunately, converting ESC’s efficiently into a specific celltype is an extremely challenging task, as we still have very little knowledge about the dynamical events that govern such transformations. In this regard, dynamically controlling the expression level of the transcription factor Nanog effectively will allow us to develop better methods to differentiate ESCs in a lineage-specific manner. It is known that Nanog plays a crucial role in retaining the stemness of ESCs by maintaining a robust heterogeneous expression pattern within an ESC population under various culture conditions. However, the overall noise and the proportion of intrinsic and extrinsic fluctuations in Nanog mRNA and protein expressions are mostly orchestrated by adjusting the related transcriptional events. In this context, miRNAs are well-known entities, which are often exploited to modify the number of fluctuations of their target genes both at the mRNA and protein levels. This suggests that the stringent Nanog expression heterogeneity can be eventually controlled by introducing Nanog transcript specific miRNAs in ESCs. Employing a stochastic mathematical modeling approach (**SFig.2**), in this work, by analyzing a relatively simple Nanog transcriptional regulatory network (**Fig. 1** and **SFig. 1**), we highlighted the possible ways to fine-tune the Nanog expression heterogeneity by altering either the binding efficiency of miR-296 and Nanog transcripts or the miR-296 expression level.

Experimentally, Schmiedel et al. [27] recently engineered a set of miRNAs to manipulate the expression heterogeneity of specific genes and measured both intrinsic and extrinsic fluctuations at the protein level. This exhaustive study revealed that the miRNAs mostly reduce the intrinsic noise as a function of total protein expression with increasing binding efficiency of miRNA-mRNA complexes. The extrinsic fluctuation mostly increases under the same situation. However, total noise initially decreases and later increases with a rise in total protein number, but has a higher influence of intrinsic fluctuation. Our stochastic simulation study unravels that heterogeneity related to Nanog expression can be altered in a varied manner (**Fig. 2–5**) depending on how we engineer and express the miR-296. We can have scenarios similar to the observation made by Schmiedel et al. [27] (**Fig. 2A-B**), or can come up with a completely different fluctuation pattern (**Fig. 2C-D**), where intrinsic fluctuation and even the total noise escalate with an increase in the transcript specific binding efficiency of miR-296-Nanog transcript. It is noteworthy to mention that in the case of Nanog, under normal WT-type and different inhibitory conditions, the extent of extrinsic noise is reportedly higher than the intrinsic noise. However, as the binding efficiency between miR-296-Nanog transcript rises, the effective drop (**Fig. 2A-B** and **SFig. 3**) or rise (**Fig. 2C-D** and **SFig. 3**) in intrinsic noise dominates the overall noise. This is how Nanog expression heterogeneity (**SFig. 3**) is distinctly different than the observations made by Schmiedel et al. [27].

Our stochastic simulations further demonstrate that nonspecific miR-296 will lead to even more complicated fluctuation patterns (**Fig. 3–5**) related to Nanog expression. Our stochastic simulations predict that depending on how the Nanog transcripts get degraded form the miR-296-mRNA complexes, we can have situations where (i) the intrinsic and extrinsic noise vary in a completely contrasting manner (**Fig. 3A** and **Fig. 3C**) as a function of specific binding efficiency of miR-296 and Nanog transcript, (ii) under high binding efficiency of miR-296 and Nanog mRNA, the intrinsic and extrinsic fluctuations pass through a minimum and maximum (**Fig. 4B**), respectively, as a function of total mRNA level under any specific expression level of miR-296, and (iii) the extrinsic fluctuation can even decrease (**Fig. 5C-D**) as a function of total transcript number in presence of various inhibitory culture conditions. Importantly, we have provided explanations for most of our observations qualitatively made from our stochastic simulations. Moreover, extensive sensitivity analysis (**Fig. 6A-C**) revealed how to alter the Nanog heterogeneity in a particular manner (**Fig. 6D-F**) to modify the intrinsic:extrinsic ratio from the experimentally observed value of 45:55 [11]. Interestingly, the model can manifest the bi-stable nature of Nanog regulation (**Fig. 6G-H**), and can even qualitatively describe the lethality effect [63] of 2i condition on Nanog-negative cells (**Fig. 6I**). Thus, our simulation results put forward new possibilities to maneuver Nanog expression heterogeneity in a contextdependent manner.

To summarize, our proposed stochastic model essentially indicates that by simply altering miR-296 regulation in a particular manner, it is possible to modify Nanog expression heterogeneity systematically. It is important to note that a significant portion of our model simulations in the presence of miR-296, at this point is a set of thought experiments that remains to be validated experimentally. However, most of our model predictions can be challenged experimentally by developing miR-296 having different complementarity with the respective Nanog transcripts. Experimentally, by genetically transforming the 3’UTR region of the Nanog transcripts (MS2 and PP7) either identically (for nonspecific binding) or in a specific manner [27] (for specific binding), we can achieve the appropriate experimental conditions to test out our predictions by performing a specific set of experiments [27] to quantify heterogeneity at either mRNA or protein level of Nanog. The extent of specificity can be altered by producing small bulges (one or more than one) in the 3’UTR region of the transcript of interest [27]. Thus, we strongly believe that our modeling study provides an efficient method to devise strategies to differentiate ESC’s in a controlled way towards a specific lineage by adjusting Nanog expression heterogeneity. Moreover, our stochastic simulation method will find wide applicability in the future, as it can be implemented in a generic way to study fluctuation dynamics for any gene under the influence of miR regulation.

## Acknowledgments

Thanks are due to IIT Bombay for providing fellowship to TS.

## Reference

1 Hyslop L, Stojkovic M, Armstrong L, Walter T, Stojkovic P, Przyborski S, Herbert M, Murdoch A, Strachan T & Lako M (2005) Downregulation of NANOG Induces Differentiation of Human Embryonic Stem Cells to Extraembryonic Lineages. Stem Cells 23, 1035–1043.

2 Lie KH, Tuch BE & Sidhu KS (2012) Suppression of NANOG induces efficient differentiation of human embryonic stem cells to pancreatic endoderm. Pancreas 41, 54–64.

3 Macarthur BD, Sevilla A, Lenz M, Müller FJ, Schuldt BM, Schuppert AA, Ridden SJ, Stumpf PS, Fidalgo M, Ma’ayan A, Wang J & Lemischka IR (2012) Nanog-dependent feedback loops regulate murine embryonic stem cell heterogeneity. Nat. Cell Biol. 14, 1139–1147.

4 Faddah DA, Wang H, Cheng AW, Katz Y, Buganim Y & Jaenisch R (2013) Single-cell analysis reveals that expression of nanog is biallelic and equally variable as that of other pluripotency factors in mouse ESCs. Cell Stem Cell 13, 23–29.

5 Filipczyk A, Gkatzis K, Fu J, Hoppe PS, Lickert H, Anastassiadis K & Schroeder T (2013) Biallelic expression of nanog protein in mouse embryonic stem cells. Cell Stem Cell 13, 12–13.

6 Kalmar T, Lim C, Hayward P, Muñoz-Descalzo S, Nichols J, Garcia-Ojalvo J & Arias AM (2009) Regulated fluctuations in Nanog expression mediate cell fate decisions in embryonic stem cells. PLoS Biol. 7, 33–36.

7 Elowitz MB, Levine AJ, Siggia ED & Swain PS (2002) Stochastic gene expression in a single cell. Science (80-.). 297, 1183–1186.

8 Mcadams HH & Arkin A (1997) Stochastic mechanisms in gene expression. Proc. Natl. Acad. Sci. U. S. A. 94, 814–819.

9 Spudich Jl Fau – Koshland Jr. DE & Koshland DE J (1976) -Non-genetic individuality: chance in the single cell. Nature 262, 467–471.

10 Ochiai H, Sugawara T, Sakuma T & Yamamoto T (2014) Stochastic promoter activation affects Nanog expression variability in mouse embryonic stem cells. Sci. Rep. 4, 1–9.

11 Samanta T & Kar S (2019) Dynamical Reorganization of Transcriptional Events Governs Robust Nanog Heterogeneity. J. Phys. Chem. B 123, 5246–5255.

12 Ambros V (2004) The functions of animal microRNAs. Nature 431, 350–355.

13 Ehrenreich IM & Purugganan M (2008) MicroRNAs in plants: Possible contributions to phenotypic diversity. Plant Signal. Behav. 3, 829–30.

14 Shenouda SK & Alahari SK (2009) MicroRNA function in cancer: Oncogene or a tumor suppressor? Cancer Metastasis Rev. 28, 369–378.

15 Alvarez-Garcia I & Miska EA (2005) MicroRNA functions in animal development and human disease. Development 132, 4653–4662.

16 Miska EA (2005) How microRNAs control cell division, differentiation and death. Curr. Opin. Genet. Dev. 15, 563–568.

17 Fuchs Y & Steller H (2011) Programmed cell death in animal development and disease. Cell 147, 742–758.

18 Olde Loohuis NFM, Kos a., Martens GJM, Van Bokhoven H, Nadif Kasri N & Aschrafi a. (2012) MicroRNA networks direct neuronal development and plasticity. Cell. Mol. Life Sci. 69, 89–102.

19 Baehrecke EH (2003) miRNAs: Micro managers of programmed cell death. Curr. Biol. 13, 473–475.

20 Schratt G (2009) MicroRNAs at the synapse. Nat. Rev. Neurosci. 10, 842–849.

21 Lozano C, Duroux-Richard I, Firat H, Schordan E & Apparailly F (2019) MicroRNAs: Key regulators to understand osteoclast differentiation? Front. Immunol. 10, 1–13.

22 Hafner M, Landthaler M, Burger L, Khorshid M, Berninger P, Rothballer A, Jr MA, Munschauer M, Ulrich A, Wardle GS, Dewell S, Zavolan M & Tuschl T (2010) Transcritpome wide identification of RNA binding protein and microRNA target sites by PAR-CLIP. Cell 141, 129–141.

23 Zeng Y, Yi R & Cullen BR (2003) MicroRNAs and small interfering RNAs can inhibit mRNA expression by similar mechanisms. Proc. Natl. Acad. Sci. 100, 9779–9784.

24 Bhattacharyya SN, Habermacher R, Martine U, Closs EI & Filipowicz W (2006) Relief of microRNA-Mediated Translational Repression in Human Cells Subjected to Stress. Cell 125, 1111–1124.

25 Rana TM. (2007) Illuminating the silence: understanding the structure and function of small RNAs. Mol. Cell. Biol. 8, 23–36.

26 Ryan B, Joilin G & Williams JM (2015) Plasticity-related microRNA and their potential contribution to the maintenance of long-term potentiation. Front. Mol. Neurosci. 8, 1–17.

27 Schmiedel JM, Klemm SL, Zheng, Y, Sahay A, Bluthgen N, Marks DS, & Oudenaarden A (2015) MicroRNA control of protein expression noise. SCIENCE. 6230, 128–132.

28 Tay Y, Zhang J, Thomson AM, Lim B & Rigoutsos I (2008) MicroRNAs to Nanog, Oct4 and Sox2 coding regions modulate embryonic stem cell differentiation. Nature 455, 1124–1128.

29 Chickarmane V, Troein C, Nuber UA, Sauro HM & Peterson C (2006) Transcriptional dynamics of the embryonic stem cell switch. PLoS Comput. Biol. 2, 1080–1092.

30 Chikarmane V & Peterson C (2008) A computational model for understanding stem cell, trophectoderm and endoderm lineage determination. PLoS One 3, 1–8.

31 Chickarmane V, Olariu V & Peterson C (2012) Probing the role of stochasticity in a model of the embryonic stem cell – heterogeneous gene expression and reprogramming efficiency. BMC Sy st. Biol. 6.

32 Akberdin IR, Omelyanchuk NA, Fadeev SI, Leskova NE, Oschepkova EA, Kazantsev F V., Matushkin YG, Afonnikov DA & Kolchanov NA (2018) Pluripotency gene network dynamics: System views from parametric analysis. PLoS One 13, 1–24.

33 Gillespie DT (1976) A general method for numerically simulating the stochastic time evolution of coupled chemical reactions. J. Comput. Phys. 22, 403–434.

34 Huh D & Paulsson J (2011) Random partitioning of molecules at cell division. Proc. Natl. Acad. Sci. U. S. A. 108, 15004–15009.

35 Dictenberg J (2012) Genetic encoding of fluorescent RNA ensures a bright future for visualizing nucleic acid dynamics. Trends Biotechnol. 30, 621–626.

36 Guet CC, Bruneaux L, Min TL, Siegal-gaskins D, Figueroa I, Emonet T & Cluzel P (2008) Minimally invasive determination of mRNA concentration in single living bacteria. Nucleic Acids Res. 36.

37 Singer ZS, Yong J, Tischler J, Hackett JA, Altinok A, Surani MA, Cai L & Elowitz MB (2014) Dynamic Heterogeneity and DNA Methylation in Embryonic Stem Cells. Mol. Cell 55, 319–331.

38 Munsky B (2012) Using Gene Expression Noise to understand Gene regulation. Science 336, 183–187.

39 Nicolas D, Phillips NE & Naef F (2017) What shapes eukaryotic transcriptional bursting? Mol. Biosyst. 13, 1280–1290.

40 Abranches E, Bekman E & Henrique D (2013) Generation and Characterization of a Novel Mouse Embryonic Stem Cell Line with a Dynamic Reporter of Nanog Expression. PLoS One 8, 1–13.

41 Navarro P, Festuccia N, Colby D, Gagliardi A, Mullin NP, Zhang W, Karwacki-Neisius V, Osorno R, Kelly D, Robertson M & Chambers I (2012) OCT4/SOX2-independent Nanog autorepression modulates heterogeneous Nanog gene expression in mouse ES cells. EMBO J. 31, 4547–4562.

42 Karwacki-Neisius V, Göke J, Osorno R, Halbritter F, Ng JH, Weiße AY, Wong FCK, Gagliardi A, Mullin NP, Festuccia N, Colby D, Tomlinson SR, Ng HH & Chambers I (2013) Reduced Oct4 expression directs a robust pluripotent state with distinct signaling activity and increased enhancer occupancy by Oct4 and Nanog. Cell Stem Cell 12, 531–545.

43 Newman JRS, Ghaemmaghami S, Ihmels J, Breslow DK, Noble M, DeRisi JL & Weissman JS (2006) Single-cell proteomic analysis of S. cerevisiae reveals the architecture of biological noise. Nature 441, 840–846.

44 Wang J, Levasseur DN & Orkin SH (2008) Requirement of Nanog dimerization for stem cell self-renewal and pluripotency. Proc Natl Acad Sci U S A 105, 6326–6331.

45 Bain J, Plater L, Elliott M, Shpiro N, Hastie CJ, Mclauchlan H, Klevernic I, Arthur JSC, Alessi DR & Cohen P (2007) The selectivity of protein kinase inhibitors: a further update. Biochem. J. 408, 297–315.

46 Sebolt-Leopold JS & Herrera R (2004) Targeting the mitogen-activated protein kinase cascade to treat cancer. Nat. Rev. Cancer 4, 937–947.

47 Miyanari Y & Torres-Padilla ME (2012) Control of ground-state pluripotency by allelic regulation of Nanog. Nature 483, 470–473.

48 Hamazaki T, Kehoe SM, Nakano T & Terada N (2006) The Grb2/Mek Pathway Represses Nanog in Murine Embryonic Stem Cells. Mol. Cell. Biol. 26, 7539–7549.

49 Storm MP, Bone HK, Beck CG, Bourillot PY, Schreiber V, Damiano T, Nelson A, Savatier P & Welham MJ (2007) Regulation of nanog expression by phosphoinositide 3-kinase-dependent signaling in murine embryonic stem cells. J. Biol. Chem. 282, 6265–6273.

50 Ying QL, Wray J, Nichols J, Batlle-morera L, Doble B, Woodgett J, Cohen P, Smith A & Ln O (2008) The ground state of embryonic stem cell self-renewal. Nature 453, 519–523.

51 Murray JT, Campbell DG, Morrice N, Auld GC, Shpiro N, Marquez R, Peggie M, Bain J, Bloomberg GB, Grahammer F, Lang F, Wulff P, Kuhl D & Cohen P (2004) Exploitation of KESTREL to identify NDRG family members as physiological substrates for SGK1 and GSK3. Biochem. J. 384, 477–488.

52 Herbert KM, Pimienta G, DeGregorio SJ, Alexandrov A & Steitz JA (2013) Phosphorylation of DGCR8 Increases Its Intracellular Stability and Induces a Progrowth miRNA Profile. Cell Rep. 5, 1070–1081.

53 Ai Z, Shao J, Shi X, Yu M, Wu Y, Du J, Zhang Y & Guo Z (2016) Maintenance of self renewal and pluripotency in J1 mouse embryonic stem cells through regulating transcription factor and microRNA expression induced by PD0325901. Stem Cells Int. 2016.

54 Tang X, Li M, Tucker L & Ramratnam B (2011) Glycogen synthase kinase 3 beta (GSK3β) phosphorylates the RNAase III enzyme Drosha at S300 and S302. PLoS One 6, 1–6.

55 Wu Y, Liu F, Liu Y, Liu X, Ai Z, Guo Z & Zhang Y (2015) GSK3 inhibitors CHIR99021 and 6-bromoindirubin-3’-oxime inhibit microRNA maturation in mouse embryonic stem cells. Sci. Rep. 5, 22–24.

56 Kim MO, Kim SH, Cho YY, Nadas J, Jeong CH, Yao K, Kim DJ, Yu DH, Keum YS, Lee KY, Huang Z, Bode AM & Dong Z (2012) ERK1 and ERK2 regulate embryonic stem cell self-renewal through phosphorylation of Klf4. Nat. Struct. Mol. Biol. 19, 283–290.

57 Skinner SO, Xu H, Nagarkar-Jaiswal S, Freire PR, Zwaka TP & Golding I (2016) Single-cell analysis of transcription kinetics across the cell cycle. Elife 5, 1–24.

58 Zopf CJ, Quinn K, Zeidman J & Maheshri N (2013) Cell-Cycle Dependence of Transcription Dominates Noise in Gene Expression. PLoS Comput. Biol. 9, 1–12.

59 Dadiani M, Van Dijk D, Segal B, Field Y, Ben-Artzi G, Raveh-Sadka T, Levo M, Kaplow I, Weinberger A & Segal E (2013) Two DNA-encoded strategies for increasing expression with opposing effects on promoter dynamics and transcriptional noise. Genome Res. 23, 966–976.

60 Hastreiter S, Skylaki S, Loeffler D, Reimann A, Hilsenbeck O, Hoppe PS, Coutu DL, Kokkaliaris KD, Schwarzfischer M, Anastassiadis K, Theis FJ & Schroeder T (2018) Inductive and Selective Effects of GSK3 and MEK Inhibition on Nanog Heterogeneity in Embryonic Stem Cells. Stem Cell Reports 11, 58–69.

